# Characterizing the molecular and metabolic mechanisms of insecticide resistance in *Anopheles gambiae* s.l. in Faranah, Guinea

**DOI:** 10.1101/610998

**Authors:** Caleb Stica, Claire L. Jeffries, Seth R. Irish, Yaya Barry, Denka Camara, Ismael Yansane, Mojca Kristan, Thomas Walker, Louisa A. Messenger

## Abstract

**Background:** In recent years, the scale-up of long-lasting insecticidal nets (LLINs) and indoor residual spraying (IRS) has greatly reduced malaria transmission. However, malaria remains a global public health concern with the majority of disease burden in sub-Saharan Africa. Insecticide resistance is a growing problem among *Anopheles* vector populations, with potential implications for the continued effectiveness of available control interventions. Improved understanding of current resistance levels and underlying mechanisms is essential to design appropriate management strategies and to mitigate future selection for resistance.

**Methods:** *Anopheles gambiae* s.l. mosquitoes were collected from three villages in Faranah Prefecture, Guinea and their levels of susceptibility to seven insecticides were measured using CDC resistance intensity bioassays. Synergist assays with piperonyl butoxide (PBO) were also undertaken to assess the role of elevated mixed-function oxidases in resistance. RNA was extracted from 563 individuals and PCR was performed on cDNA to determine vector species, presence of target site mutations (L1014F *kdr*, N1575Y and G119S *Ace-1*), *Plasmodium falciparum* infection, and relative expression of three metabolic genes (*CYP6M2, CYP6P3* and *GSTD3*).

**Results:** In Faranah, resistance to permethrin and deltamethrin was observed, as well as possible resistance to bendiocarb. All assayed vector populations were fully susceptible to alpha-cypermethrin, pirimiphos-methyl, clothianidin and chlorfenapyr. *Plasmodium falciparum* infection was detected in 7.3% (37/508) mosquitoes tested. The L1014F *kdr* mutation was found in 100% of a sub-sample of 60 mosquitoes, supporting its fixation in the region. The N1575Y mutation was identified in 20% (113/561) of individuals, with ongoing selection evidenced by significant deviations from Hardy-Weinberg equilibrium. The G119S *Ace-1* mutation was detected in 62.1% (18/29) of mosquitoes tested and was highly predictive of bendiocarb bioassay survival. The metabolic resistance genes, *CYP6M2, CYP6P3* and *GSTD3*, were found to be overexpressed in wild resistant and susceptible *An. gambiae* s.s. populations, compared to a susceptible G3 colony. Furthermore, *CYP6P3* was significantly overexpressed in bendiocarb survivors, implicating its potential role in carbamate resistance in Faranah.

**Conclusions:** Identification of intense resistance to permethrin and deltamethrin in Faranah, is of concern, as the Guinea National Malaria Control Program (NMCP) relies exclusively on the distribution of pyrethroid-treated LLINs for vector control. Study findings will be used to guide current and future control strategies in the region.

## Background

Despite impressive progress made towards the control and elimination of malaria, this disease remains the leading cause of morbidity and mortality in the tropics, where it is estimated to have resulted in the deaths of approximately 435,000 individuals in 2017 [1]. Between 2010 and 2017, global malaria incidence has fallen by approximately 18% globally (72 to 59 cases per 1000 at risk) and by 20% in the World Health Organization African region [1] which still bears the greatest disease burden [2]. Malaria deaths have been decreasing annually, largely due to the scale-up of long-lasting insecticidal nets (LLINs) [3] and implementation of indoor residual spraying (IRS) [2]. However, progress has stalled in some areas, with an increase of 2 million cases from 2016 to 2017 [1].

Long-term intensive insecticide use to control agricultural pests and disease vectors has resulted in the selection of resistance in many insect species [4]. Beginning with the widespread use of dichlorodiphenyltrichloroethane (DDT) in the 1950-1960s, followed by the recent increase in distribution of LLINs impregnated with pyrethroids, and the broad use of the same insecticides in the agricultural industry have led to the development of resistance in mosquito populations worldwide [5]. This resistance poses a major threat to malaria control [4] where vector control is reliant primarily on insecticide-based interventions [6]. Of the 80 malaria endemic countries for which data are available from 2010 onwards, 68 countries detected decreased susceptibility to at least one insecticide among vector populations, with 57 countries reporting resistance to two or more chemical classes [1].

In Guinea, malaria remains one of the most significant diseases of public health importance, with 92% of infections caused by *Plasmodium falciparum* [7]. The national malaria prevalence is approximately 15%, reaching up to 25% in Faranah Prefecture [8]. Guinea’s tropical climate allows for year-round malaria transmission, with peak transmission from July through to October in most areas. The Guinean national vector control strategy focuses almost exclusively on the distribution of LLINs, with IRS occurring in only 1.7% of households, primarily those of workers engaged in mining operations [9]. The United States President’s Malaria Initiative (PMI) estimates that intervention coverage remains low with only 48% of households owning at least one insecticide-treated net (ITN) per every two members [9]. Given the reliance on LLINs for malaria control, the detection of nationwide pyrethroid resistance is of concern [10]. In order to safeguard malaria control efforts in the country, we characterized current insecticide resistance levels and underlying mechanisms, to inform the design of appropriate management strategies and mitigate future selection for resistance.

## Methods

### Study sites

Human landing catches (HLCs) were conducted at three sites in Faranah Prefecture (Figure 1); The villages of Balayani (10.1325, −10.7443), Foulaya (10.144633, −10.749717), and Tindo (9.9612230, −10.7016560) were selected, due to their high malaria prevalence, and were visited every other day until a sufficient sample size for testing was acquired. Following consent from the household owner, two to three fieldworkers, positioned outside of the house, collected mosquitoes landing and attempting to feed on their exposed legs and feet from 1800h to 0700h. Mosquitoes were transported back to the insectary in Faranah and provided with 10% sucrose solution prior to bioassay testing.

**Figure 1.**
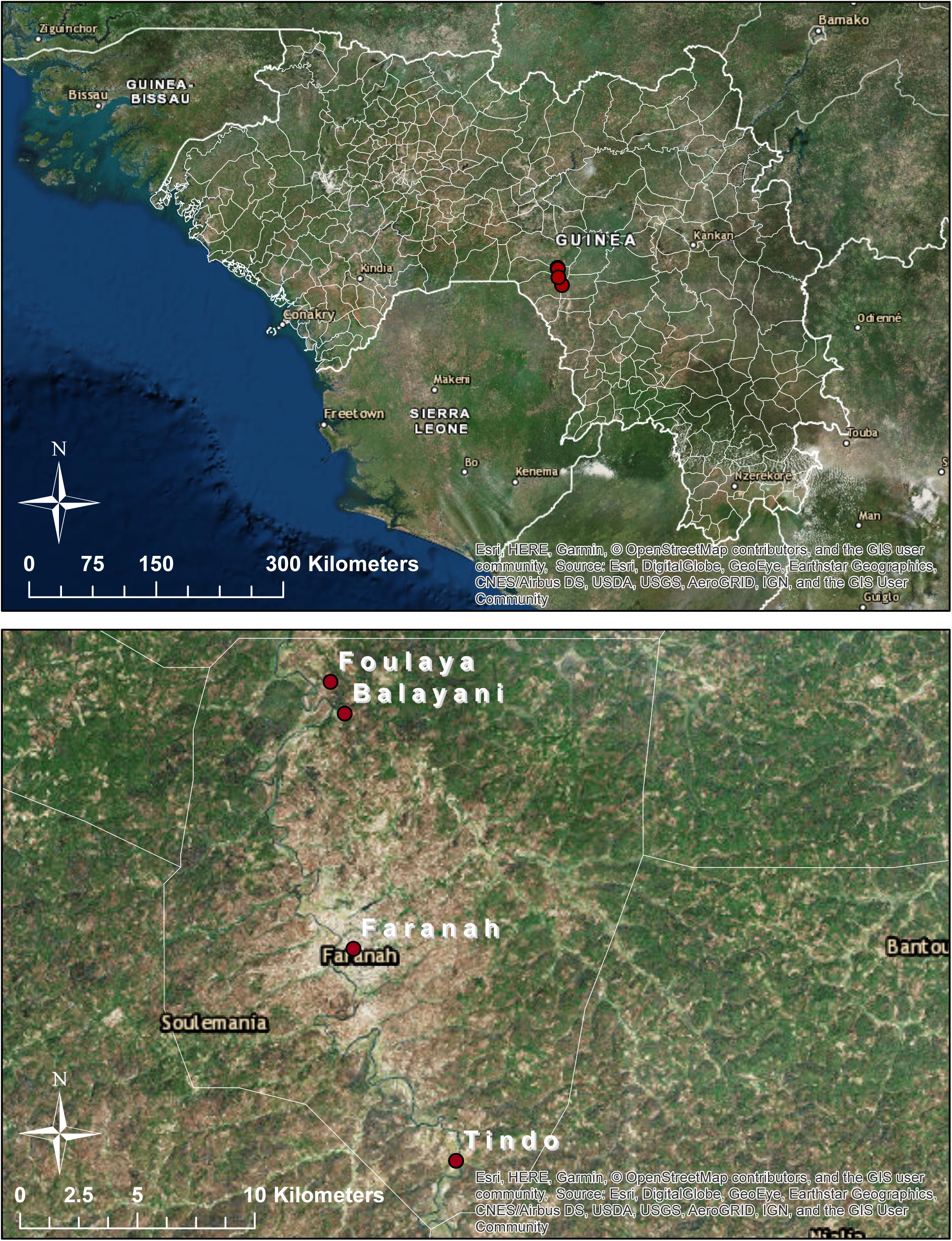
Map of study sites in Faranah Prefecture, Guinea. Human landing catches (HLCs) were performed in the villages of Balayani, Foulaya and Tindo to collect adult *An. gambiae* s.l. Larval collections were undertaken at multiple sites in the city of Faranah.

Larval collections were also performed in Faranah (10.042423, −10.740980), at sites selected through active searching and/or known to have been productive in previous years. Larvae and pupae were collected using larval dippers, ladles, buckets, and pans. Collections were then transported back to the insectary in Faranah where they were fed on ground fish food (TetraMin^®^ Tropical Fish Food Flakes). Pupae were removed on a daily basis, placed in plastic cups, and transferred to mosquito cages. Emerging adult mosquitoes were provided 10% sucrose solution prior to bioassay testing. All mosquito samples were collected between 25^th^ June and 20^th^ July 2018, at the start of the long rainy season.

### Susceptibility tests

CDC bottle bioassays were performed using 250 ml Wheaton bottles with adult female mosquitoes of varying ages caught in HLCs from the three villages (each village was tested separately), or 2-5-day old female mosquitoes raised from larvae in the insectary. Bioassays were performed at the Centre de Santé Marché in Faranah; mosquitoes were held in the insectary for no more than 48 hours prior to testing. Following CDC guidelines, bottles were coated with alpha-cypermethrin (12.5 μg/bottle), deltamethrin (12.5 μg/bottle), permethrin (21.5 μg/bottle), and bendiocarb (12.5 μg/bottle) at 1, 2, 5, and 10 times the diagnostic dose, and with pirimiphos-methyl (20 μg/bottle), clothianidin (90 μg/bottle), and chlorfenapyr (100 μg/bottle) at the diagnostic dose. Stock solutions for all insecticides and synergists were prepared using 95-98% ethanol as a solvent for all, clothianidin also included 58.8 μg Mero (Sigma-Aldrich, USA), to circumvent issues previously experienced with bottle coating and insecticide crystallization. Approximately 20-25 mosquitoes were introduced into each assay bottle and mortality was recorded in all bottles at the start of the assay and at 15-minute intervals for up to 30 minutes (for alpha-cypermethrin, deltamethrin, permethrin, bendiocarb, and pirimiphos-methyl) or 60 minutes (for clothianidin and chlorfenapyr). Mosquitoes exposed to chlorfenapyr were held for an additional 24h in paper cups, with access to 10% sugar solution. In each bioassay, a control bottle, coated with 98% ethanol, was run in parallel. Mortality was defined as the inability of a mosquito to stand or fly in a coordinated manner [11]. Synergist CDC bottle bioassays were performed using piperonyl butoxide (PBO) to investigate the potential role of detoxifying enzyme families in resistance. Bottles were coated with 100 µg of PBO and mosquitoes were exposed for 60 minutes, followed by exposure to pyrethroid treated bottles. Adult mosquitoes reared from larvae were exposed for 30 minutes to permethrin and deltamethrin for comparison to wild caught adults. At the end of each exposure period, separate individual surviving (resistant), knocked down or dead (susceptible), and control mosquitoes were dipped in ethanol and then preserved in RNAlater® (Thermo Fisher Scientific, UK) at 4°C in Faranah, for a maximum of three weeks, and subsequently at −80°C for downstream molecular analyses at the London School of Hygiene and Tropical Medicine (LSHTM).

### RNA extraction

Single mosquitoes were homogenized using a Qiagen Tissue Lyser II (Qiagen, Hilden, Germany) with 3mm stainless steel beads and RNA was extracted using Qiagen 96 RNeasy Kits according to the manufacturer’s instructions (Qiagen, Hilden, Germany). RNA was eluted in 45 μL of RNase-free water and stored at −70°C. Specimens were selected from all three study villages, including representative susceptible and resistant individuals per insecticide per concentration. A High Capacity cDNA Reverse Transcription kit (Applied Biosystems) was used to perform reverse transcription on eluted RNA. Reactions were performed in a Bio-Rad T100™ Thermal Cycler which cycled for 10 minutes at 25°C, 120 minutes at 37°C, and 5 minutes at 85°C. The resulting cDNA was then stored at −20°C.

### Identification of *Anopheles gambiae* species complex

Mosquitoes were morphologically identified as *Anopheles gambiae* s.l. in the field prior to CDC bottle bioassay testing. A sub-set of 480 samples were further identified using an end point PCR assay developed by Santolamazza, *et al*. [12]. This assay amplifies the SINE200 insertion, a highly repetitive ∼200 bp element which is widespread in the *An. gambiae* s.s genome [12]. Samples were prepared with forward (5’-TCGCCTTAGACCTTGCGTTA-3’) and reverse (5’-CGCTTCAAGAATTCGAGATAC-3’) primers, and amplifications performed in 20 µL reactions containing 2 µL cDNA, 2 µL of each primer (10 µM), 4 µL H_2_O, and 10 µL 2X Hot Start Taq PCR Master Mix (New England Biolabs). The cycling conditions in the Bio-Rad T100™ Thermal Cycler were: 10 minutes at 94°C; 35 cycles of 30 seconds at 94°C, 30 seconds at 54°C, and 60 seconds at 72°C; then 10 minutes at 72°C. Products were run on 2% agarose gels in an Invitrogen E-gel iBase™ Real-Time Transilluminator. Amplification products of 479 bp or 249 bp were considered indicative of *An. coluzzii* or *An. gambiae* s.s., respectively; both bands indicate a hybrid individual. The *An. gambiae* s.s. form has an identical banding pattern to that of *An. melas* and *An. quadriannulatus* (249 bp), however, due to the geographical location of sampling, it is highly unlikely that these species were present.

### *Plasmodium falciparum* detection

A total of 508 whole body mosquito samples (collected in HLCs) were tested for the presence of *P. falciparum* using a real-time assay targeting the cytochrome c oxidase subunit 1 (*cox1*) mitochondrial gene of *P. falciparum* according to Boissière *et al.* [13]. This assay is highly sensitive and specific, capable of detecting the target gene at all stages of the *P. falciparum* lifecycle [13]. Samples were prepared with forward (5’-TTACATCAGGAATGTTATTGC-3’) and reverse (5’-ATATTGGATCTCCTGCAAAT-3’) primers, and amplifications performed in 10 µL reactions containing 2 µL cDNA, 1 µL of each primer (10 µM), 1 µL H_2_O, and 5 µL2X Roche FastStart Essential DNA green master mix containing SYBR Green. Samples were run on a Roche Lightcycler^®^ 96 Real-Time PCR system for 15 minutes at 95°C, followed by 35 cycles of 15 seconds at 95°C and 30 seconds at 58°C. Fluorescence results were analyzed using Roche Lightcycler^®^ 96 software. Positive controls were used from gDNA extracted from a cultured *P. falciparum*-infected blood sample (parasitaemia of ∼10%) in addition to the inclusion of no template controls (NTCs).

### Characterization of resistance mutations: **target site mutations**

A sub-sample of 60 mosquitoes was randomly selected to be tested for West African L1014F *kdr* mutation, given the high frequency of this allele and its fixation in many parts of Guinea and West Africa [10, 14]. The PCR master mix was prepared according to MR4 guidelines [15]. Primers IPCF (5’-GATAATGTGGATAGATTCCCCGACCATG-3’), AltRev (5’-TGCCGTTGGTGCAGACAAGGATG-3’), WEST WT (5’-GGTCCATGTTAATTTGCATTACTTACGAATA-3’), and West:West (5’-CTTGGCCACTGTAGTGATAGGAAATGTT-3’) were used to detect the L1014F allele. Amplifications were performed in 25 µL reaction containing 2 µL cDNA, 1 µL IPCF (2.5 pmol/µL), 1 µL AltRev (2.5 pmol/µL), 1 µL West WT (25.0 pmol/µL), 3 µL West:West (8.0 pmol/µL), 4.5 µL H_2_O, and 12.5 µL 2X Hot Start Taq PCR Master Mix (New England Biolabs). Samples were run on a Bio-Rad T100™ Thermal Cycler and cycled for 5 minutes at 95°C, followed by 35 cycles of 30 seconds at 95°C, 30 seconds at 59°C, and 30 seconds at 72°C, and a final extension step of 5 minutes at 72°C. PCR products were separated on 2% agarose gels in an Invitrogen E-gel iBase™ Real-Time Transilluminator. A control band at 314 bp indicated a successful reaction, a band at 214 bp indicated the susceptible wild type, and a band at 156 bp indicated the resistance mutation.

Detection of the N1575Y mutation was carried out on 570 samples (including the 60 individuals who tested positive for L1014F *kdr*), using a TaqMan PCR assay developed by Jones *et al.* [16]. Forward primer (5’-TGGATCGCTAGAAATGTTCATGACA-3’), reverse primer (5’-CGAGGAATTGCCTTTAGAGGTTTCT-3’), and two probes: Yprobe (5’-TTTTTCATTGCATAATAGTAC-3’) and Nprobe (5’-ATTTTTTTCATTGCATTATAGTAC-3’) were used to detect the presence of the wildtype and the mutation allele. HEX and FAM fluorophores were used due to their different excitation wavelengths, ensuring no interference: excitation of HEX showed no mutation, while excitation of HEX and FAM at similar Ct values indicated the N1575Y mutation. 20 µL reactions containing 2 µL cDNA, 1 µL each primer (10 µM), 0.5 µL each probe, 5 µL H_2_O, and 10 µL QuantiTect Probe Master Mix were prepared in plates and run on an Agilent Technologies Stratagene Mx3005P qPCR system and cycled according to the Quantitect(™) Probe PCR Handbook guidelines (15 minutes at 95°C; 35 cycles of 15 seconds at 95°C and 60 seconds at 60°C). Positive controls from gDNA extracted from known *An. gambiae* s.s. with the N1575Y mutation and without the mutation were included on each run in addition to no template controls (NTCs).

A subsample of 30 mosquitoes which were resistant or susceptible to bendiocarb were tested for the presence of the G119S *Ace-1* mutation using a TaqMan PCR assay, according to Weill *et al*. [17]. Samples were prepared with degenerate primers Moustdir1 (5′-CCGGGNGCSACYATGTGGAA-3′) and Moustrev1 (5′-ACGATMACGTTCTCYTCCGA-3’), and amplifications performed in 20 µL reactions containing 2 µL cDNA, 2 µL each primer (10 µM), 4 µL H_2_O, and 10 µL 2X Hot Start Taq PCR Master Mix (New England Biolabs). Samples were loaded into a Biorad T100™ Thermal Cycler for 3 minutes at 95°C, followed by 30 cycles of 30 seconds at 95°C, 30 seconds at 52°C, and 1 minute at 72°C, and a final step of 10 minutes at 72°C. The resulting PCR fragments were then digested with *Alu*I restriction enzyme (Thermo Scientific) for 16 hours, according to the manufacturer’s instructions, and run on 2% agarose gels in an Invitrogen E-gel iBase™ Real-Time Transilluminator. 194 bp undigested PCR products indicated the susceptible allele and 74 bp and 120 bp digested fragments indicated the presence of the resistant allele. Presence of all three product sizes indicated the sample was heterozygous. Positive controls from gDNA extracted from known *An. gambiae s.s*. that were homozygous susceptible (SS), homozygous resistant (RR) and heterozygous individuals (RS) for G119S *Ace-1* were included in addition to no template controls (NTCs).

### Characterization of resistance mechanisms: metabolic gene expression

The relative gene expression of two cytochrome-dependent monooxygenases: *CYP6P3, CYP6M2*, and glutathione s-transferase, *GSTD3*, was analyzed in 461 individuals from Guinea and 41 susceptible G3 individuals from a colony at LSHTM, using quantitative qPCR relative to the housekeeping gene *RPS7*, according to Yahouédo *et al.* [18]. These genes were selected based upon their significant overexpression in other neighboring West African vector populations [18, 19]. Each gene used the following primers: *RPS7* forward (5’-ATTGCCGAGCGCCGCATTCT-3’) and reverse (5’-GACGCGGATACGCTTGCCGA-3’) primers, *CYP6M2* forward (5’-TCGGGATGTGTGCGTTCGGC-3’) and reverse (5’-TCGTGTCTCGCACCGCGTTC-3’) primers, *CYP6P3* forward (5’-TGTGATTGACGAAACCCTTCGGAAG-3’) and reverse (5’-ATAGTCCACAGACGGTACGCGGG-3’) primers, and *GSTD3* forward (5’-CTAAGCTTAATCCGCAACATACCA-3’) and reverse (5’-GTGTCATCCTTGCCGTACAC-3’) primers. For each gene, 10 µL reactions were prepared containing 2 µL cDNA, 1 µL each primer (10 µM), 1 µL H_2_O, and 5 µL 2X Roche FastStart Essential DNA green master mix containing SYBR Green. Prepared reactions were loaded into the Agilent Technologies Stratagene Mx3005P qPCR system which cycled for 10 minutes at 95°C; 35 cycles of 10 seconds at 95°C, 22 seconds at 60°C, and 10 seconds at 72°C; followed by a melt curve. Serial dilutions were performed on selected samples for each of the four genes and relative standard curves produced using the Stratagene MxPro qPCR software (Agilent Technologies). Using the same software, sample Ct values could then be used to generate relative quantities, accounting for each assays’ efficiency, and the expression level of each metabolic gene tested could be normalised to the housekeeping gene *RPS7*.

### Data analysis

Data were recorded on pre-prepared data sheets and entered into an excel spreadsheet. Control mortality in bioassays never exceeded 5%, thus correction using Abbott’s formula was not necessary. Mosquito mortality was analyzed according to WHO criteria: 98-100% mortality at 30 minutes of exposure indicates ‘susceptibility’, 90-97% mortality suggests ‘possible resistance’ and <90% indicates the presence of ‘resistance’ [4]. GraphPad Prism 7 (GraphPad Software) was used for statistical analysis (t-tests, Fisher’s exact tests and chi-squared tests). Microsoft Excel was used to calculate proportions and construct resistance graphs. Stratagene MxPro qPCR software (Agilent Technologies) was used to produce relative standard curves for genotypic analysis.

## Results

### Mosquito collections and species identification

A total of 2,597 female *An. gambiae* s.l. mosquitoes were either wild-caught using HLCs from three villages in Faranah Prefecture (Balayani= 956, Foulaya=914, and Tindo=589) or raised from larvae collected in Faranah (n=138). Of those, 480 were selected for molecular species identification, with 466 (97.1%) determined to be *An. gambiae* s.s., 6 (1.3%) identified as *An. coluzzii*, and 6 (1.3%) identified as hybrids; results were inconclusive for 2 (0.4%) individuals.

### Insecticide resistance intensity

Levels of susceptibility to seven insecticides (permethrin, deltamethrin, alpha-cypermethrin, bendiocarb, pirimiphos-methyl, clothianidin and chlorfenapyr) were assessed across three villages in Faranah (Balayani, Foulaya, and Tindo). Overall, each study site displayed comparable resistance profiles for deltamethrin (pyrethroid) and bendiocarb (carbamate); no significant association between mosquito mortality and sample site was observed (χ^2^=6.495, *p*=0.0899 for deltamethrin 1X; χ^2^=1.338, *p*=0.5122, χ =2.38, *p*=0.304 and χ^2^=0.903, *p*=0.637 for bendiocarb at 1X, 2X and 5X, respectively). However, mosquito mortality following permethrin (pyrethroid) exposure varied among villages in Faranah (χ^2^=8.573, *p*=0.035 and χ^2^=29.58, *p*<0.0001, for 1X and 2X, respectively).

Considering HLC-collected adult mosquitoes from Faranah Prefecture as a whole, resistance was consistently observed to permethrin and deltamethrin. Permethrin gave the lowest mortality of all insecticides tested with 4% [95% CI 1%, 11%] and 15% [95% CI 9%, 24%] mosquito mortality at 1X and 2X the diagnostic doses, respectively (Figure 2 and Table 1). Resistance to deltamethrin was also evident, but to a lesser degree; average mosquito mortality to the diagnostic dose was 86% [95% CI 77%, 91%] (Figure 2). Possible resistance was observed to bendiocarb, with mortality ranging between 94-97% at 1X, 2X, and 5X concentrations. Mosquitoes were found to be susceptible to the diagnostic doses of all other insecticides with mortalities >98% for alpha-cypermethrin, chlorfenapyr, clothianidin, and pirimiphos-methyl (Figure 2 and Table 1).

**Table 1.**
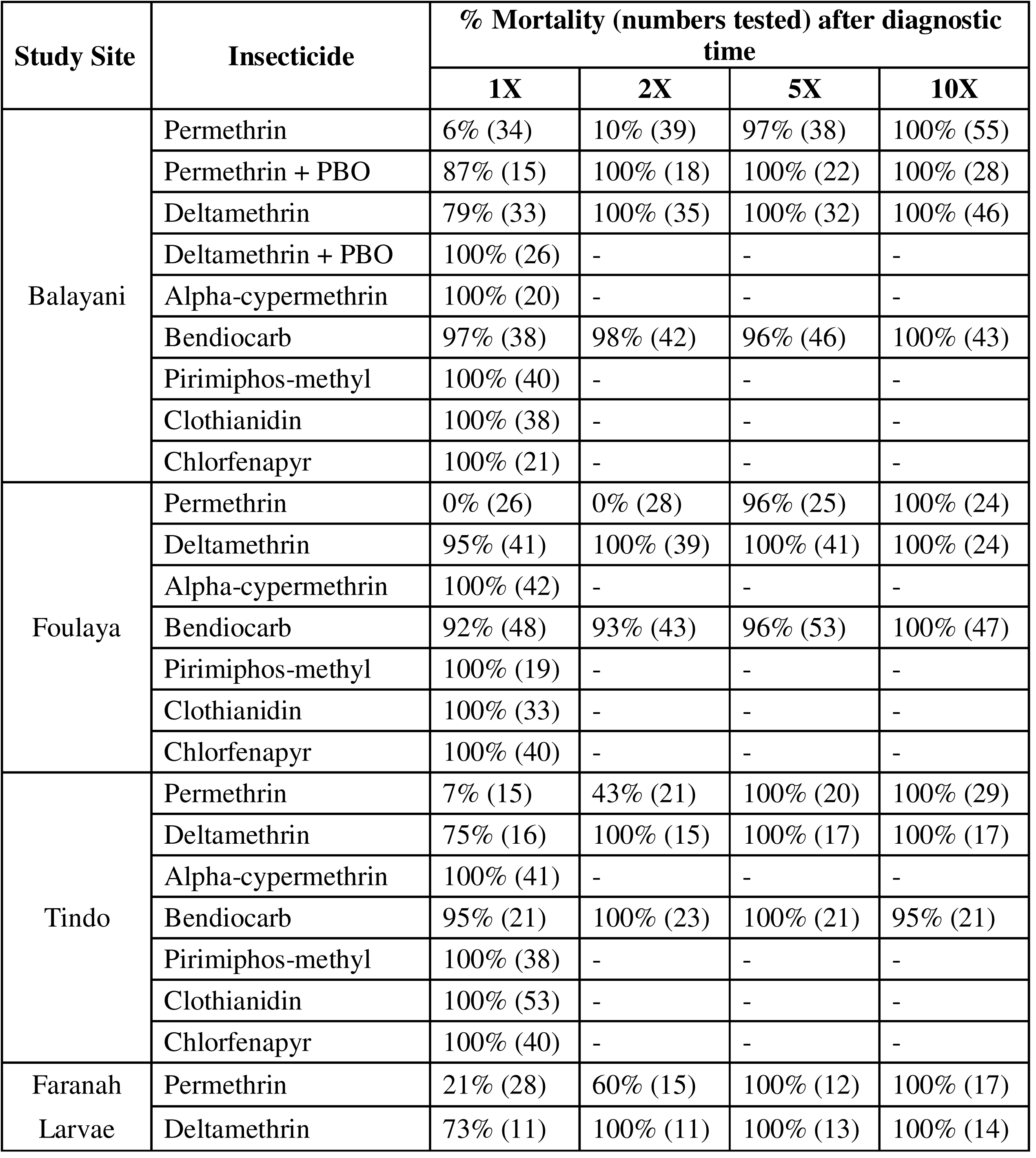
Percentage mortality (and numbers tested) of *Anopheles gambiae* s.l. in CDC resistance intensity assays conducted at four sites in Faranah Prefecture, Guinea.

| Study Site | Insecticide | % Mortality (numbers tested) after diagnostic time |  |  |  |
| --- | --- | --- | --- | --- | --- |
|  |  | 1X | 2X | 5X | 10X |
| Balayani | Permethrin | 6% (34) | 10% (39) | 97% (38) | 100% (55) |
|  | Permethrin + PBO | 87% (15) | 100% (18) | 100% (22) | 100% (28) |
|  | Deltamethrin | 79% (33) | 100% (35) | 100% (32) | 100% (46) |
|  | Deltamethrin + PBO | 100% (26) | - | - | - |
|  | Alpha-cypermethrin | 100% (20) | - | - | - |
|  | Bendiocarb | 97% (38) | 98% (42) | 96% (46) | 100% (43) |
|  | Pirimiphos-methyl | 100% (40) | - | - | - |
|  | Clothianidin | 100% (38) | - | - | - |
|  | Chlorfenapyr | 100% (21) | - | - | - |
| Foulaya | Permethrin | 0% (26) | 0% (28) | 96% (25) | 100% (24) |
|  | Deltamethrin | 95% (41) | 100% (39) | 100% (41) | 100% (24) |
|  | Alpha-cypermethrin | 100% (42) | - | - | - |
|  | Bendiocarb | 92% (48) | 93% (43) | 96% (53) | 100% (47) |
|  | Pirimiphos-methyl | 100% (19) | - | - | - |
|  | Clothianidin | 100% (33) | - | - | - |
|  | Chlorfenapyr | 100% (40) | - | - | - |
| Tindo | Permethrin | 7% (15) | 43% (21) | 100% (20) | 100% (29) |
|  | Deltamethrin | 75% (16) | 100% (15) | 100% (17) | 100% (17) |
|  | Alpha-cypermethrin | 100% (41) | - | - | - |
|  | Bendiocarb | 95% (21) | 100% (23) | 100% (21) | 95% (21) |
|  | Pirimiphos-methyl | 100% (38) | - | - | - |
|  | Clothianidin | 100% (53) | - | - | - |
|  | Chlorfenapyr | 100% (40) | - | - | - |
| Faranah Larvae | Permethrin | 21% (28) | 60% (15) | 100% (12) | 100% (17) |
|  | Deltamethrin | 73% (11) | 100% (11) | 100% (13) | 100% (14) |

**Figure 2.**
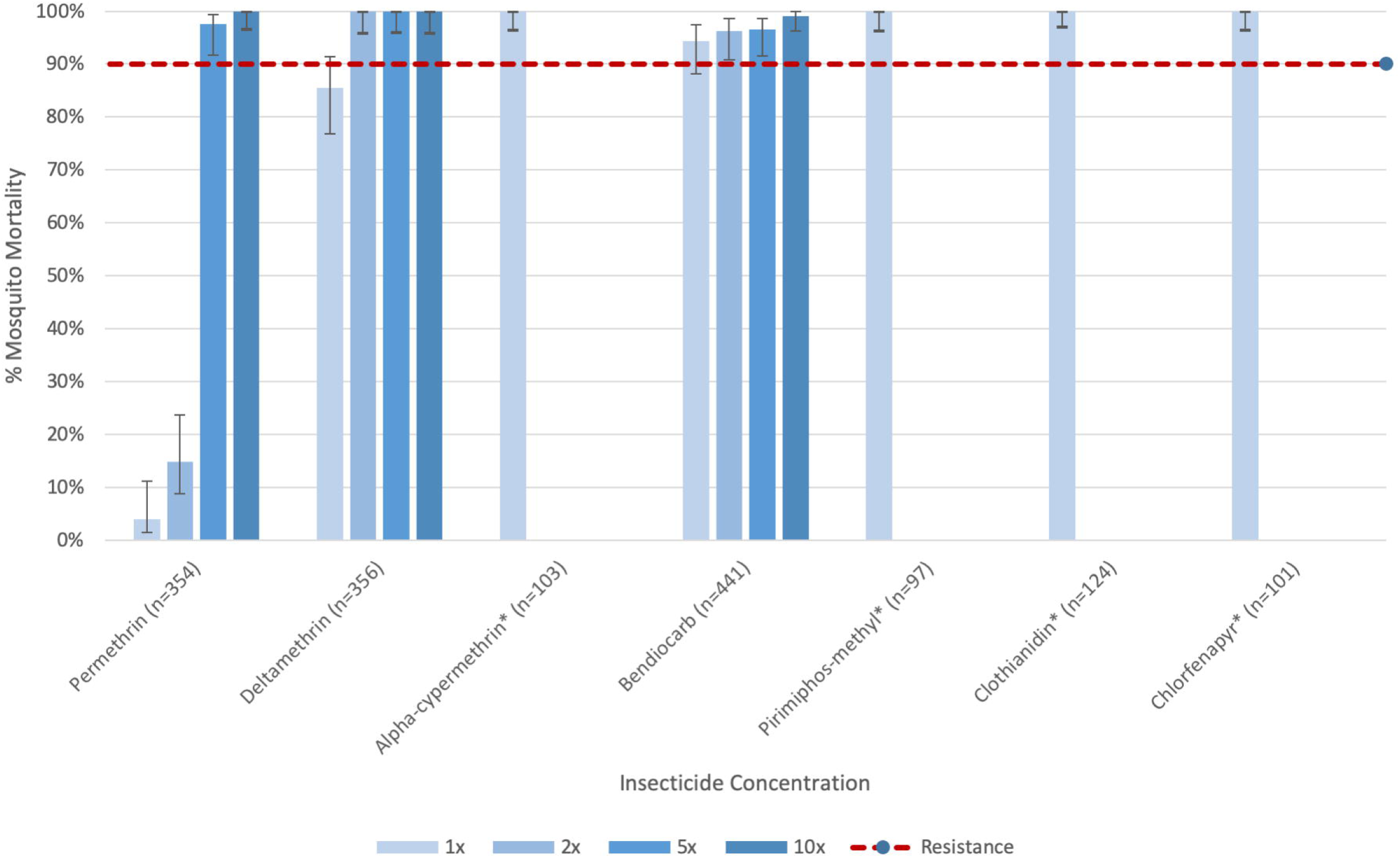
Pooled CDC resistance intensity assay data for all tested insecticides (permethrin, deltamethrin, alpha-cypermethrin, bendiocarb, pirimiphos-methyl, clothianidin and chlorfenapyr) in Faranah Prefecture, Guinea for adult wild-caught mosquitoes. Mortality below 90% indicates the presence of confirmed resistance. *only tested at diagnostic dose.

Minor variations in levels of resistance were apparent between study villages. The most intense permethrin resistance was measured in Foulaya, with average mosquito mortality of 0% [95% CI 0%, 13%] following exposure to either 1X or 2X the diagnostic dose, relative to 6% [95% CI 2%, 19%] and 10% [95% CI 4%, 24%] in Balayani and 7% [95% CI 1%, 30%] and 43% [95% CI 24%, 63%] in Tindo, respectively (Table 1). Possible resistance to permethrin 5X was identified in Foulaya and Balayani (average mosquito mortality of 96% [95% CI 81%, 99%] and 97% [95% CI 87%, 100%], respectively). By comparison, the lowest levels of resistance to the diagnostic dose of deltamethrin were observed in Foulaya, with average mosquito mortality of 95% [95% CI 84%, 99%], compared to 79% [95% CI 62%, 89%] and 75% [95% CI 51%, 90%] in Balayani and Tindo, respectively.

Two to five-day old mosquitoes raised in the insectary from larvae demonstrated moderately similar resistance profiles to locally collected adult populations of varying physiological age (Table 1). Both permethrin and deltamethrin resistance was present, with average mosquito mortality of 21% [95% CI 10%, 40%] and 60% [95% CI 36%, 80%] for permethrin 1X and 2X doses, respectively, and 73% [95% CI 43%, 90%] mortality for deltamethrin 1X. Comparing resistance profiles in larvae to that observed in each village, a statistical difference was found for permethrin 1X in Foulaya (*χ*^2^=6.014, *p*=0.0142) and in all villages combined at 1X (*χ*^2^=7.407, *p*=0.0065); permethrin 2X in Balayani (*χ*^2^=14.62, *p*=0.0001) and Foulaya (*χ*^2^=20.753, *p*<0.0001), and all villages combined at 2X (*χ*^2^=15.2, *p*=0.0001); and deltamethrin 1X in Foulaya (*χ*^2^=4.72, *p*=0.0298). Reared mosquitoes were found to be susceptible to all other concentrations of permethrin and deltamethrin with mortalities of 100%.

Synergist bioassays were performed on a sub-sample of pyrethroid resistant mosquitoes from Balayani. Pre-exposure to PBO and subsequent permethrin or deltamethrin treatment resulted in partial or complete abolishment of resistance. Mortality to permethrin increased from 6% to 87% at 1X, and from 10% to 100% at 2X; mortality following deltamethrin exposure increased from 79% to 100% at 1X.

### *Plasmodium falciparum* detection

Of the 508 mosquitoes tested for the presence of *P. falciparum*, 37 individuals were positive (indicating the presence of any parasite lifecycle stage), giving an infection rate of 7.3%. There was no significant difference in *P. falciparum* infection among study villages (χ^2^=1.358, *p*=0.507). Considering pooled insecticide results (permethrin, deltamethrin, and bendiocarb), susceptible mosquitoes were more likely to be infected with *P*. *falciparum* (7.6%; 33/432) than resistant mosquitoes (2.8%; 4/144) (χ^2^=4.236, *p*=0.039). However, no statistical association between survival to insecticide exposure and *P. falciparum* infection was observed per individual chemical (Fisher’s exact: permethrin 2X *p*=0.37, 5X *p*=0.54; deltamethrin 1X *p*=0.43; bendiocarb 1X *p*=0.58, 2X *p*=0.73, 5X *p=*0.6 and 10X *p=*0.54)

### Characterization of resistance mechanisms: target site mutations

Of the sub-sample of 60 mosquitoes which were tested for the L1014F *kdr* allele, this mutation was identified in all *An. gambiae* s.s. samples (100%). Of the 570 individuals which were assayed for the presence of N1575Y, this mutation was detected in 20% (113/561; 9 specimens failed to amplify) (Table 2). Four (0.7%) individuals were heterozygous for this mutation, while the remainder were homozygous. Across Faranah, the frequency of N1575Y did not vary significantly (*χ*^2^=4.4819, *p*=0.214 for comparisons between Balayani, Foulaya, Tindo, and larvae from Faranah). In addition, no significant association was observed between the presence of the N1575Y mutation and the ability to survive exposure to pyrethroid insecticides (permethrin 1X χ^2^ =0.015, *p*=0.902; 2X χ^2^ =0.189, *p*=0.664; 5X χ^2^ =0.814, *p*=0.359; deltamethrin 1X χ^2^ =0.934, *p*=0.334). Significant deviations from Hardy-Weinberg equilibrium were observed in almost all phenotyped specimens, indicative of ongoing selection (Table 2).

**Table 2.**
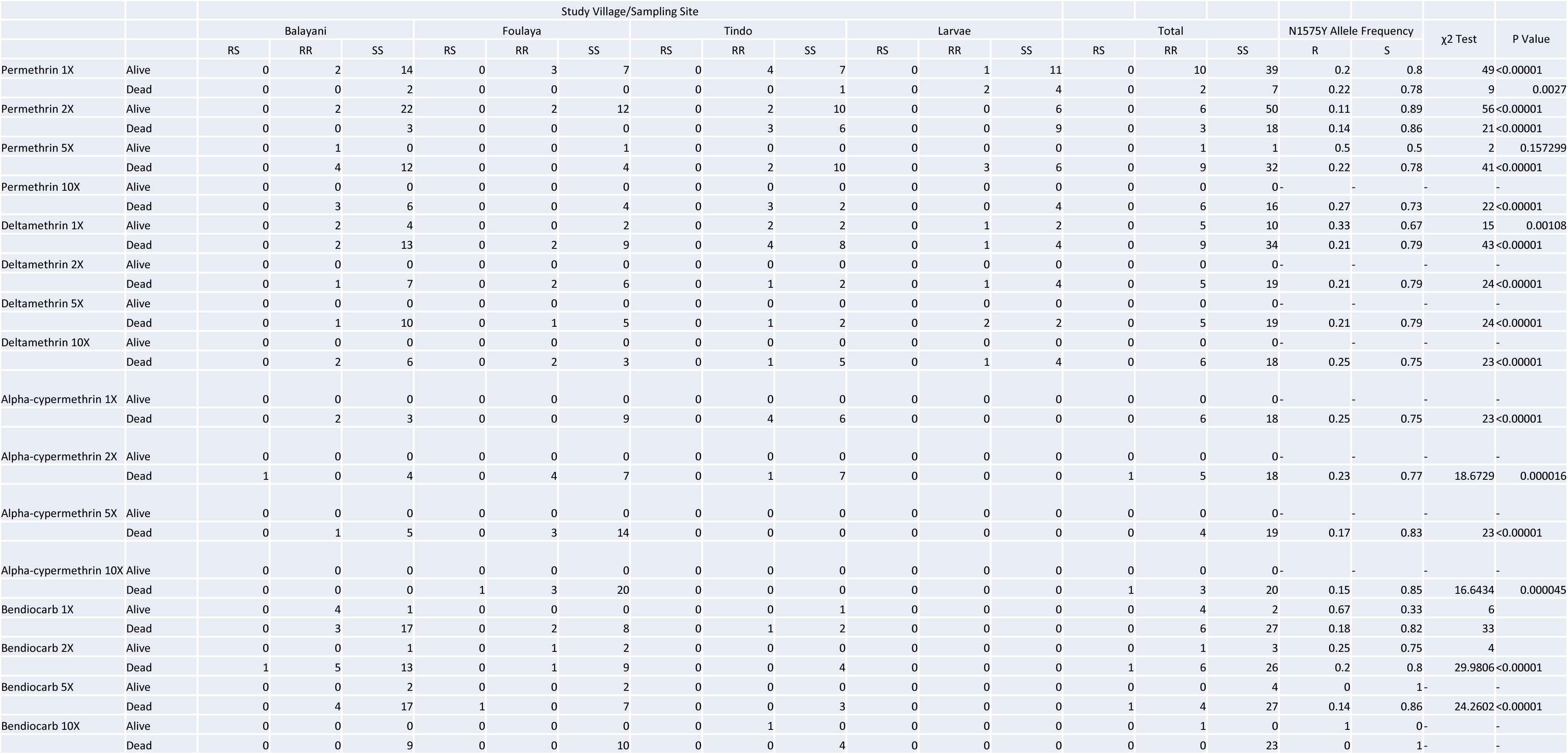
N1575Y allele frequencies and *P* values for chi-square tests for deviations from Hardy-Weinberg equilibrium in *An. gambiae* s.s. from four sites in Faranah Prefecture, Guinea.

|  |  | Study Village/Sampling Site |  |  |  |  |  |  |  |  |  |  |  |  |  |  |  |  |  |  |
| --- | --- | --- | --- | --- | --- | --- | --- | --- | --- | --- | --- | --- | --- | --- | --- | --- | --- | --- | --- | --- |
|  |  | Balayani |  |  | Foulaya |  |  | Tindo |  |  | Larvae |  |  | Total |  |  | N1575Y Allele Frequency |  | χ2 Test | P Value |
|  |  | RS | RR | SS | RS | RR | SS | RS | RR | SS | RS | RR | SS | RS | RR | SS | R | S |  |  |
| Permethrin 1X | Alive | 0 | 2 | 14 | 0 | 3 | 7 | 0 | 4 | 7 | 0 | 1 | 11 | 0 | 10 | 39 | 0.2 | 0.8 | 49 | <0.00001 |
|  | Dead | 0 | 0 | 2 | 0 | 0 | 0 | 0 | 0 | 1 | 0 | 2 | 4 | 0 | 2 | 7 | 0.22 | 0.78 | 9 | 0.0027 |
| Permethrin 2X | Alive | 0 | 2 | 22 | 0 | 2 | 12 | 0 | 2 | 10 | 0 | 0 | 6 | 0 | 6 | 50 | 0.11 | 0.89 | 56 | <0.00001 |
|  | Dead | 0 | 0 | 3 | 0 | 0 | 0 | 0 | 3 | 6 | 0 | 0 | 9 | 0 | 3 | 18 | 0.14 | 0.86 | 21 | <0.00001 |
| Permethrin 5X | Alive | 0 | 1 | 0 | 0 | 0 | 1 | 0 | 0 | 0 | 0 | 0 | 0 | 0 | 1 | 1 | 0.5 | 0.5 | 2 | 0.157299 |
|  | Dead | 0 | 4 | 12 | 0 | 0 | 4 | 0 | 2 | 10 | 0 | 3 | 6 | 0 | 9 | 32 | 0.22 | 0.78 | 41 | <0.00001 |
| Permethrin 10X | Alive | 0 | 0 | 0 | 0 | 0 | 0 | 0 | 0 | 0 | 0 | 0 | 0 | 0 | 0 | 0- | - | - | - | - |
|  | Dead | 0 | 3 | 6 | 0 | 0 | 4 | 0 | 3 | 2 | 0 | 0 | 4 | 0 | 6 | 16 | 0.27 | 0.73 | 22 | <0.00001 |
| Deltamethrin 1X | Alive | 0 | 2 | 4 | 0 | 0 | 2 | 0 | 2 | 2 | 0 | 1 | 2 | 0 | 5 | 10 | 0.33 | 0.67 | 15 | 0.00108 |
|  | Dead | 0 | 2 | 13 | 0 | 2 | 9 | 0 | 4 | 8 | 0 | 1 | 4 | 0 | 9 | 34 | 0.21 | 0.79 | 43 | <0.00001 |
| Deltamethrin 2X | Alive | 0 | 0 | 0 | 0 | 0 | 0 | 0 | 0 | 0 | 0 | 0 | 0 | 0 | 0 | 0- | - | - | - | - |
|  | Dead | 0 | 1 | 7 | 0 | 2 | 6 | 0 | 1 | 2 | 0 | 1 | 4 | 0 | 5 | 19 | 0.21 | 0.79 | 24 | <0.00001 |
| Deltamethrin 5X | Alive | 0 | 0 | 0 | 0 | 0 | 0 | 0 | 0 | 0 | 0 | 0 | 0 | 0 | 0 | 0- | - | - | - | - |
|  | Dead | 0 | 1 | 10 | 0 | 1 | 5 | 0 | 1 | 2 | 0 | 2 | 2 | 0 | 5 | 19 | 0.21 | 0.79 | 24 | <0.00001 |
| Deltamethrin 10X | Alive | 0 | 0 | 0 | 0 | 0 | 0 | 0 | 0 | 0 | 0 | 0 | 0 | 0 | 0 | 0- | - | - | - | - |
|  | Dead | 0 | 2 | 6 | 0 | 2 | 3 | 0 | 1 | 5 | 0 | 1 | 4 | 0 | 6 | 18 | 0.25 | 0.75 | 23 | <0.00001 |
| Alpha-cypermethrin 1X | Alive | 0 | 0 | 0 | 0 | 0 | 0 | 0 | 0 | 0 | 0 | 0 | 0 | 0 | 0 | 0- | - | - | - | - |
|  | Dead | 0 | 2 | 3 | 0 | 0 | 9 | 0 | 4 | 6 | 0 | 0 | 0 | 0 | 6 | 18 | 0.25 | 0.75 | 23 | <0.00001 |
| Alpha-cypermethrin 2X | Alive | 0 | 0 | 0 | 0 | 0 | 0 | 0 | 0 | 0 | 0 | 0 | 0 | 0 | 0 | 0- | - | - | - | - |
|  | Dead | 1 | 0 | 4 | 0 | 4 | 7 | 0 | 1 | 7 | 0 | 0 | 0 | 1 | 5 | 18 | 0.23 | 0.77 | 18.6729 | 0.000016 |
| Alpha-cypermethrin 5X | Alive | 0 | 0 | 0 | 0 | 0 | 0 | 0 | 0 | 0 | 0 | 0 | 0 | 0 | 0 | 0- | - | - | - | - |
|  | Dead | 0 | 1 | 5 | 0 | 3 | 14 | 0 | 0 | 0 | 0 | 0 | 0 | 0 | 4 | 19 | 0.17 | 0.83 | 23 | <0.00001 |
| Alpha-cypermethrin 10X | Alive | 0 | 0 | 0 | 0 | 0 | 0 | 0 | 0 | 0 | 0 | 0 | 0 | 0 | 0 | 0- | - | - | - | - |
|  | Dead | 0 | 0 | 0 | 1 | 3 | 20 | 0 | 0 | 0 | 0 | 0 | 0 | 1 | 3 | 20 | 0.15 | 0.85 | 16.6434 | 0.000045 |
| Bendiocarb 1X | Alive | 0 | 4 | 1 | 0 | 0 | 0 | 0 | 0 | 1 | 0 | 0 | 0 | 0 | 4 | 2 | 0.67 | 0.33 | 6 |  |
|  | Dead | 0 | 3 | 17 | 0 | 2 | 8 | 0 | 1 | 2 | 0 | 0 | 0 | 0 | 6 | 27 | 0.18 | 0.82 | 33 |  |
| Bendiocarb 2X | Alive | 0 | 0 | 1 | 0 | 1 | 2 | 0 | 0 | 0 | 0 | 0 | 0 | 0 | 1 | 3 | 0.25 | 0.75 | 4 |  |
|  | Dead | 1 | 5 | 13 | 0 | 1 | 9 | 0 | 0 | 4 | 0 | 0 | 0 | 1 | 6 | 26 | 0.2 | 0.8 | 29.9806 | <0.00001 |
| Bendiocarb 5X | Alive | 0 | 0 | 2 | 0 | 0 | 2 | 0 | 0 | 0 | 0 | 0 | 0 | 0 | 0 | 4 | 0 | 1- | - | - |
|  | Dead | 0 | 4 | 17 | 1 | 0 | 7 | 0 | 0 | 3 | 0 | 0 | 0 | 1 | 4 | 27 | 0.14 | 0.86 | 24.2602 | <0.00001 |
| Bendiocarb 10X | Alive | 0 | 0 | 0 | 0 | 0 | 0 | 0 | 1 | 0 | 0 | 0 | 0 | 0 | 1 | 0 | 1 | 0- | - | - |
|  | Dead | 0 | 0 | 9 | 0 | 0 | 10 | 0 | 0 | 4 | 0 | 0 | 0 | 0 | 0 | 23 | 0 | 1- | - | - |

Of the subsample of 30 mosquitoes which survived or died following bendiocarb exposure, the G119S *Ace-1* mutation was present in 62% (18/29 individuals; 1 specimen failed to amplify) (Table 3), with allele frequencies ranging from 0.25 to 1.0. Seventeen specimens were homozygous, the remaining individual was heterozygous. There was a statistically significant association between the presence of the G119S *Ace-1* mutation and mosquito survival after bendiocarb exposure at 1X and 2X (Fisher’s exact: *p*=0.00108 and *p*=0.0238 respectively). Significant deviations from Hardy-Weinberg equilibrium for G119S *Ace-1*, were limited to mosquitoes which died following exposure to bendiocarb 2X and 5X (Table 3).

**Table 3.**
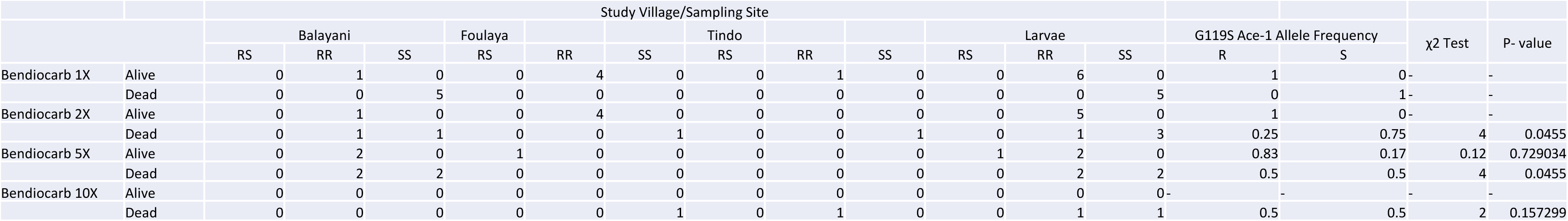
G119S *Ace-1* allele frequencies and *P* values for chi-square tests for deviations from Hardy-Weinberg equilibrium in *An. gambiae* s.s. from four sites in Faranah Prefecture, Guinea.

### Characterization of resistance mechanisms: metabolic gene expression

461 samples identified as *An. gambiae* s.s. were tested for the expression of *CYP6M2, CYP6P3*, and *GSTD3* genes relative to the housekeeping gene *RPS7* and compared to 41 *An. gambiae* s.s. samples from a susceptible G3 colony. These three genes were significantly overexpressed in the majority of wild-caught *An. gambiae* s.s. when compared to the susceptible G3 laboratory strain (Table 4 and Figure 3).

**Table 4.**
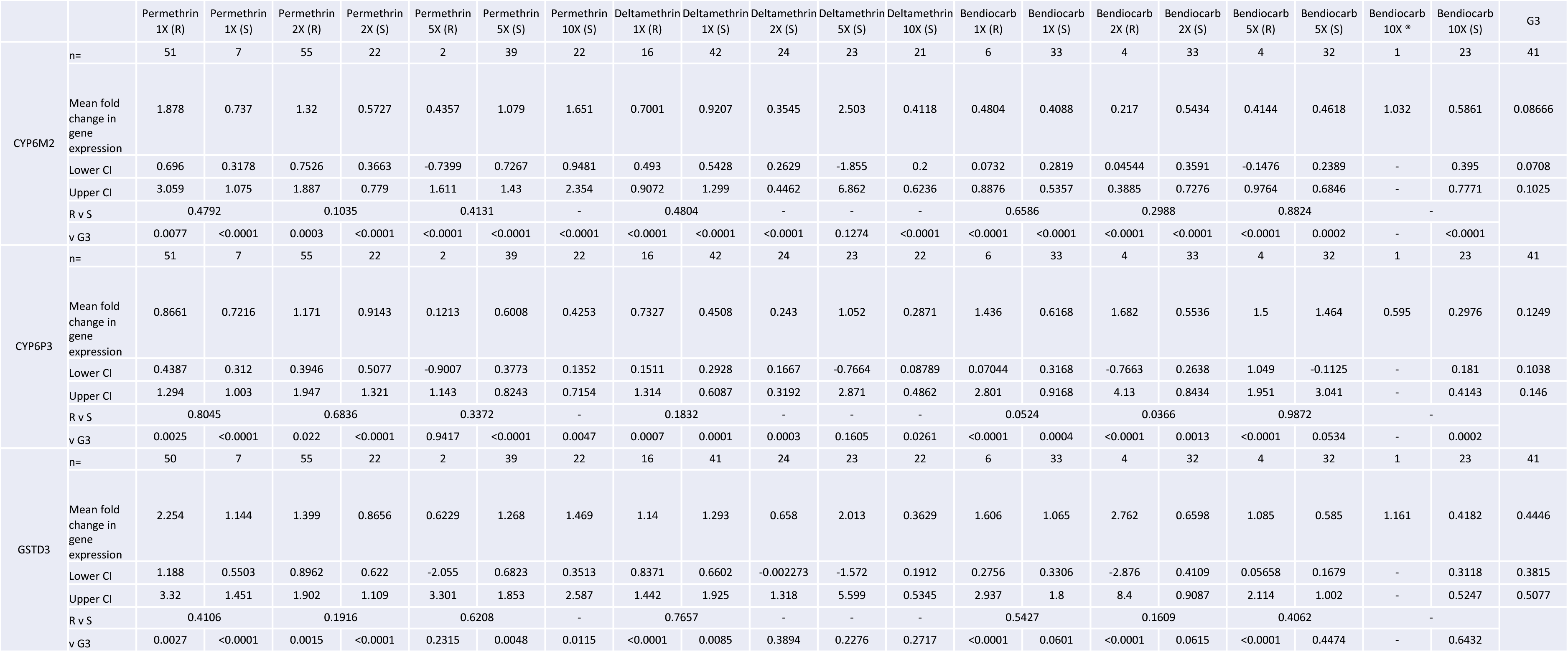
Mean fold change in relative expression of *CYP6M2, CYP6P3*, and *GSTD3*, and associated 95% confidence intervals, between *An. gambiae* s.s. populations in the Faranah Prefecture and the *An. gambiae* s.s. susceptible G3 colony. (R) = resistant or alive, (S) = susceptible or dead.

**Figure 3.**
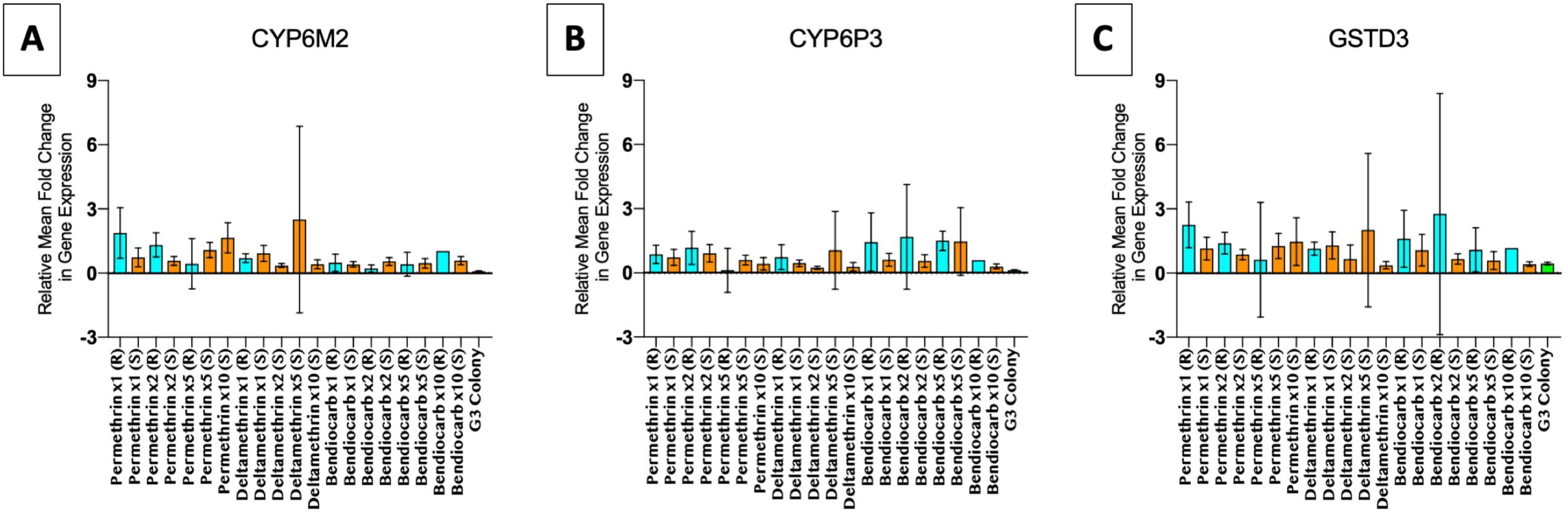
Relative expression of genes (*CYP6M2, CYP6P3*, and *GSTD3*) and associated 95% confidence intervals, for three insecticides that showed resistance at varying concentrations, normalized to the housekeeping gene *RPS7* in *An. gambiae* s.s. populations in Faranah Prefecture and in the *An. gambiae* s.s. susceptible G3 colony. (R) = resistant or alive, (S) = susceptible or dead.

The greatest changes in gene expression were observed for *GSTD3*, with an average mean fold change of 1.19 among wild *An. gambiae* s.s. compared to 0.44 in G3 colony individuals; average levels of *CYP6M2* and *CYP6P3* were 0.84 and 0.79 compared to 0.09 and 0.12 between field and colony mosquitoes, respectively. Among Guinean vectors, a significantly higher expression of *CYP6P3* was observed between individuals which survived bendiocarb exposure at 1X and 2X, compared to those that died (1.44 *vs.* 0.62; *p*=0.0524 and 1.68 *vs.* 0.55; *p*=0.0366, respectively) (Table 4). However, no significant changes in gene expression of *CYP6M2* and *GSTD3* were apparent between wild *An. gambiae* s.s. which survived or died after insecticide exposure at any dose.

## Discussion

We assessed the susceptibility of *An. gambiae* s.l. populations from Faranah, Guinea to seven public health insecticides and characterized underlying resistance mechanisms. Intense resistance to permethrin and deltamethrin was apparent, with survivors at 5X and 2X the diagnostic doses, respectively; minor heterogeneity in mosquito mortality was also evident across this restricted geographic area. By comparison, no evidence of alpha-cypermethrin resistance was observed in Faranah, with 100% mosquito mortality following exposure. Increased tolerance to particular pyrethroids in this region is not unexpected, given LLINs are the sole insecticide-based malaria control strategy implemented in Guinea. Deltamethrin LLINs were distributed nationwide in 2013 (Netprotect^®^) and 2016 (PermaNet^®^ 2.0 and Yorkool^®^) by the NMCP [10], thus a resulting increase in deltamethrin resistance would be anticipated. However, the higher levels of permethrin resistance demonstrated in this study could be the result of control efforts prior to 2013 or concurrent use of this insecticide (and/or others capable of facilitating cross-resistance) in agriculture activities. Alternatively, L1014F *kdr*, which our study indicated was fixed in Faranah in 2018, as well as in other parts of Guinea [10, 21], has been reported to contribute more to resistance to type I (permethrin) *versus* type II (alpha-cypermethrin and deltamethrin) pyrethroids [22].

In Faranah, susceptibility to all other classes of insecticides under evaluation was confirmed, excluding the carbamate, bendiocarb, which showed the development of possible resistance with mortality ranging between 94% and 97% in the region. This emerging resistance may also be attributable to carbamate use in the agricultural setting [10], as has been documented previously in other parts of Guinea [21]. The organophosphate, pirimiphos-methyl, achieved complete mosquito mortality in this study. Currently, large-scale government-funded, widespread IRS activities are not underway in Guinea. However, knowledge of susceptibility in relation to current LLIN use will help to guide potential IRS implementation in the future [9]. Local mosquito populations were also susceptible to two new insecticides under consideration for public health use, clothianidin (neonicotinoid) and chlorfenapyr (pyrrole), strengthening the evidence that net impregnations and IRS formulations with these insecticides may be successful in future vector control efforts. Likewise, absence of alpha-cypermethrin resistance supports the deployment of Interceptor G2^®^ LLINs (containing a combination of alpha-cypermethrin and chlorfenapyr) in prospective LLIN distributions.

Regarding underlying resistance mechanisms, decreased susceptibility to pyrethroids in Faranah was mediated both by target site mutations and overexpression of metabolic enzymes. The L1014F *kdr* mutation was detected in all tested samples, which is consistent with high frequencies of this mutation observed throughout the country [10, 21]. A second mutation in the voltage-gated sodium channel, N1575Y, was confirmed in 20% of vectors, with strong evidence for ongoing selection, as evidenced by significant deviations from Hardy-Weinberg allele frequencies; however this mutation was not associated with mosquito survival following insecticide exposure. A synergistic relationship between the L1014F *kdr* and N1575Y mutations has been established in other parts of West Africa, where it has been shown to enhance resistance to pyrethroids and DDT, and potentially compensate for fitness costs incurred by L1014F *kdr* [16]. By comparison, in Maferinyah, Guinea, this mutation was present at similar frequencies to Faranah and directly implicated in phenotypic resistance to permethrin [21]. The ability of the synergist PBO to re-establish full or partial susceptibility to both permethrin and deltamethrin indicates that mixed function oxidases (MFOs) play some role in the resistance reported. However, the expression of metabolic enzymes examined in this study (*CYP6M2, CYP6P3*, and *GSTD3*), while significantly upregulated in wild *An. gambiae* s.s. compared to the susceptible G3 colony, did not statistically differ between our pyrethroid resistant and susceptible wild-caught mosquitoes. Further exploratory analyses of our wild populations are warranted to characterize additional metabolic resistance pathways. Given the influence of MFOs as a predominant resistance mechanism to pyrethroids in the area and recent findings of improved protection with permethrin and PBO impregnated nets [20], the Guinea NMCP should consider including these next-generation nets in future vector control activities.

The G119S *Ace-1* mutation, which confers resistance to carbamate and organophosphate insecticides [17] was found in this study to be highly predictive of bioassay survival to bendiocarb. Furthermore, *CYP6P3* was significantly overexpressed in bendiocarb survivors, suggesting it may also be responsible for decreased tolerance to carbamates in Faranah. Similar results implicating a role for *CYP6P3* in bendiocarb metabolism and resistance have been reported from Côte d’Ivoire [23] and over-expression of this P450 has been documented among many multi-insecticide resistant field populations [24-28].

In this study, mosquitoes that were reared from larvae collected from sites in the town of Faranah showed a higher mortality rate to permethrin at 1X and 2X the diagnostic dose, when compared to the wild-caught adult population of varying physiological age. This finding contradicts previous studies which have suggested that field-collected adults have higher mortality rates due to the mixture of ages and blood-feeding statuses in these populations [29], and the reported decline of phenotypic resistance with increasing mosquito age [30]. This observation strengthens the importance of utilizing the same mosquito collection method and assay preparation method when testing field-collected mosquitoes [29] and cautions against the use of such data interchangeably.

Malaria prevalence in Faranah Prefecture is approximately 15%, reaching up to 25% [8]; mosquito infection rates in this study were 7.3%. Because whole mosquito bodies were screened for *P. falciparum* by PCR, it is also possible that, in addition to infection or infectivity, a positive sample could be indicative of a mosquito which recently fed upon an infected human bloodmeal. More importantly, insecticide susceptible vectors were more likely to be infected with *P. falciparum*, but this phenomenon could not be ascribed to a single insecticide or resistance mechanism,.

Additional considerations for the interpretation of study data include that in the tested population of *An. gambiae* s.l., *An. gambiae* s.s. was the predominant species at 97.1%, with *An. coluzzii* representing 1.3%, and the hybrid form 1.3%. Only two samples failed to produce any result on the *An. gambiae* complex end-point PCR assay. *An. gambiae* s.s. breed in temporary, rain-dependent breeding sites [32], which could account for its predominance during the rainy season. Future collections at multiple timepoints, utilizing different collection methods would be necessary to determine the principal species in the area. Sampling took place in only three villages in one area of Guinea, and therefore, these findings cannot be extrapolated across the entire country. However, this information will contribute to an understanding of resistance in this region, which will be valuable information for the National Malaria Control Program (NMCP). [33].

## Conclusions

Resistance to the pyrethroids, permethrin and deltamethrin, as well as possible resistance to bendiocarb were observed in Faranah Prefecture in Guinea. Both target site mutations (L1014F *kdr*, N1575Y and G119S *Ace-1*) and metabolic mechanisms of resistance are present in this field population. The complete susceptibility of vectors to alpha-cypermethrin, clothianidin, chlorfenapyr, and pirimiphos-methyl is encouraging, and these insecticides should be considered by the NMCP in future control efforts in the form of new LLIN and IRS interventions as they become approved and readily available.

## Declarations

### Ethics approval and consent to participate

The study protocol was reviewed and approved by the Comité National d’Ethique pour la Recherche en Santé (030/CNERS/17) and the institutional review boards (IRB) of the London School of Hygiene and Tropical Medicine (#14990). The protocol was reviewed at the Centers for Disease Control and Prevention, USA and determined to be non-human subject research (reference number 2018-086); all study procedures were performed in accordance with relevant guidelines and regulations. Fieldworkers participating in human landing catches were provided with malaria prophylaxis for the duration of the study.

### Consent for publication

Not applicable

### Availability of data and material

Not applicable

### Authors’ Contributions

LAM, CS, SRI, CLJ and TW designed the study and were responsible for data analysis and interpretation. CS, YB, DC, IY and MK led the entomology field activities and participated in data collection. TW, CS and CLJ performed the molecular assays. CS, LAM, TW and SRI drafted the manuscript, which was revised by co-authors. All authors read and approved the final manuscript.

## Acknowledgements

The authors would like to thank all mosquito collectors for their dedicated work.

## Disclaimer

The findings and conclusions in this report are those of the author(s) and do not necessarily represent the official position of the Centers for Disease Control and Prevention.

